# Less diverse but uniquely composed: Canopy cover and elevation shape Neotropical Arcellinida assemblages in the Peruvian Andes

**DOI:** 10.64898/2026.09.30.755618

**Authors:** Nura ElKhouri-Vidarte, Emilio Cano, Italo Treviño-Zevallos, Enrique Lara

## Abstract

The Peruvian Andes, and especially its slopes towards the Amazonian Basin are well-known biodiversity hotspots for plants and animals, which represent the visible part of biodiversity. However, not much is known about microeukaryotic diversity, despite the significant role protists play in the ecosystem. Based on a molecular survey of the diversity of Arcellinida (a group of amoebae well known for its environmental sensitivity and restricted geographical distributions), we estimated its diversity along an elevation gradient spanning more than 3000 meters. Surprisingly, Neotropical forest Arcellinida were less abundant and diverse than in most other extratropical biomes; nevertheless, communities differed strongly in their taxonomic composition as compared with forests in higher latitudes. Notably, sequences related to *Trigonopyxis* were most abundant, whereas this group occurs only marginally in extratropical settings. Elevation influenced beta diversity, showing different assembly processes occurring at different elevation belts. Canopy cover positively influenced both phylogenetic and taxonomic diversity, showing the importance of above and below ground integration in tropical forests.

## Introduction

The Neotropical region is well acknowledged for its immense diversity, characterised by a variety of ecoregions and endemisms (Antonelli & Sanmartín, 2011; Hanson et al., 2009; Raven et al., 2020). Moreover, mountains are known refuges for biodiversity and host a quarter of all species on Earth while occupying only 25% of land area (Rahbek et al., 2019). For all these reasons, the Andean mountain range is a particularly important hotspot for species richness and endemicity (Myers et al., 2000; Pennington et al., 2010). Besides its importance in species diversification (Antonelli & Sanmartín, 2011), the Andes are essential to climate regulation because of their role as a carbon sink (Chambers et al., 2001; Phillips et al., 1998) and provide ecosystem services to millions of people (Pérez-Escobar et al., 2018). A significant part of its productivity is attributed to plant growth, biomass accumulation, and diversity, which rely on the vast richness of the ecosystems. Notably, the Neotropical Andes contain approximately 10% of the world’s plant species despite covering little more than 0.5% of Earth’s surface (Mittermeier et al., 2004).

While macroorganisms have historically been the most studied in these hotspots, protists represent the bulk of eukaryotic diversity at a global level (Burki et al., 2021; Griffith, 2012) and notably, in soil where they play essential roles in the ecosystems (Geisen et al., 2018). Understanding protist diversity patterns, and how they are linked to animal and plant diversity has been a major challenge for diversity research these last decades (González-Miguéns et al., 2026a; Lara et al., 2016; Mahé et al., 2016; Pinseel et al., 2024). The immense variety of sizes, lifestyles and phylogenetic origins of protists increasingly encourages researchers to focus diversity investigations to single groups, both for consistency reasons and for practical considerations (Lara et al ., 2022).

Among the protists present in soil ecosystems, Arcellinida stands out as a very diverse clade of testate amoebae that have a characteristic self-constructed shell that, paired to molecular analyses, is used for taxonomic identification (Ribeiro et al., 2023). These organisms are known to follow macroscopic organisms diversity patterns, including geographic structuring (González-Miguéns et al., 2026b; Singer et al., 2019) and macroecological patterns (ElKhouri-Vidarte et al., 2025). Arcellinida is a model group in environmental monitoring and the study of diversity patterns since they are key players in the ecosystem cycles and functioning, as well as bioindicators (Useros et al., 2025). Indeed, they regulate the ecosystems by actively participating in nutrient cycling and trophic webs, as they can be considered great predators of the microworld (Dumack et al., 2024; Geisen et al., 2018), feeding notably on other eukaryotes (Gilbert et al., 2003, Geisen et al., 2015). Moreover, they are highly sensitive to changes in environmental conditions, making them excellent indicators of the environmental health (Soler-Zamora et al., 2021). The main drivers modulating Arcellinida communities are temperature, humidity, vegetation, and soil conditions such as pH and soil moisture (Booth, 2001; Fernández et al., 2016; Tsyganov et al., 2022). Vegetation, in particular, has been demonstrated to be a major driver for beta diversity in an elevation gradient in subtropical forests in Japan, explaining over 12% of the total variation (Tsyganov et al., 2022). The influence of climatic factors on Arcellinida diversity has been studied especially in elevation gradients. While assessing species richness, some studies have found no correlation between diversity and elevation (Mitchell, 2004), while others have identified positive, negative, and non-linear relationships (Heger et al., 2016; Krashevska et al., 2007; Tsyganov et al., 2013). Local climatic conditions are most likely responsible for diversity fluctuations along the gradients.

However, these studies often relied on morphological identification, or DNA barcoding, which although essential, are economically limiting and time-consuming techniques. Approaches like next-generation sequencing (NGS) and environmental DNA (eDNA) have only recently been used for Arcellinida (González-Miguéns et al., 2023), and they offer the potential to evaluate Arcellinida communities and diversity patterns across multiple samples simultaneously, thus overcoming these limitations (ElKhouri-Vidarte et al., 2026).

Understanding the relationship between Arcellinida diversity and environmental changes in a key ecosystem such as the Andes might allow us to infer changes in the systems and focus conservation efforts on the most urgent matters. The current situation in the Andes of Peru is concerning due to the rapid disappearance of neotropical mountain forests caused by local fires and deforestation (Intendencia Nacional de Bomberos del Perú, 2025; Koelle & Soto, 2025). Given the ecological significance of these ecosystems, it is urgent to establish a robust baseline on their diversity and the variables that modulate it.

In this context, this study aims to (i) address the knowledge gap regarding Arcellinida in the Neotropical mountains by identifying the main groups present in the ecosystems and (ii) determine the main drivers that influence Arcellinida diversity in neotropical forests. As this region is a biodiversity hotspot, we expect to find a high diversity of Arcellinida. Based on earlier results (e.g. Tsyganov et al., 2022), we anticipate that vegetation should play a key role as a diversity driver.

## Material and Methods

### Area ofstudy and sampling

Sampling was conducted in both Regions of Cusco and Madre de Dios, in Southern Peru in the buffer zone between the Manu National Park and the Amarakaeri Communal Reserve (Figure 1). Along an elevation gradient going from 969 m to 3999 m, we collected 200 soil samples following the sampling protocol of ElKhouri-Vidarte et al. (2026), but pressing the soil litter to obtain the interstitial water instead of using a mesh. All of the samples were immediately fixed after collection with LifeGuard™ Soil Preservation Solution (Qiagen), following the instructions provided by the manufacturer. A duplicate for each sample was deposited at the Museo Sur Peruano (MSP), affiliated to the Instituto Científico Michael Owen Dillon (IMOD).

**Figure 1:**
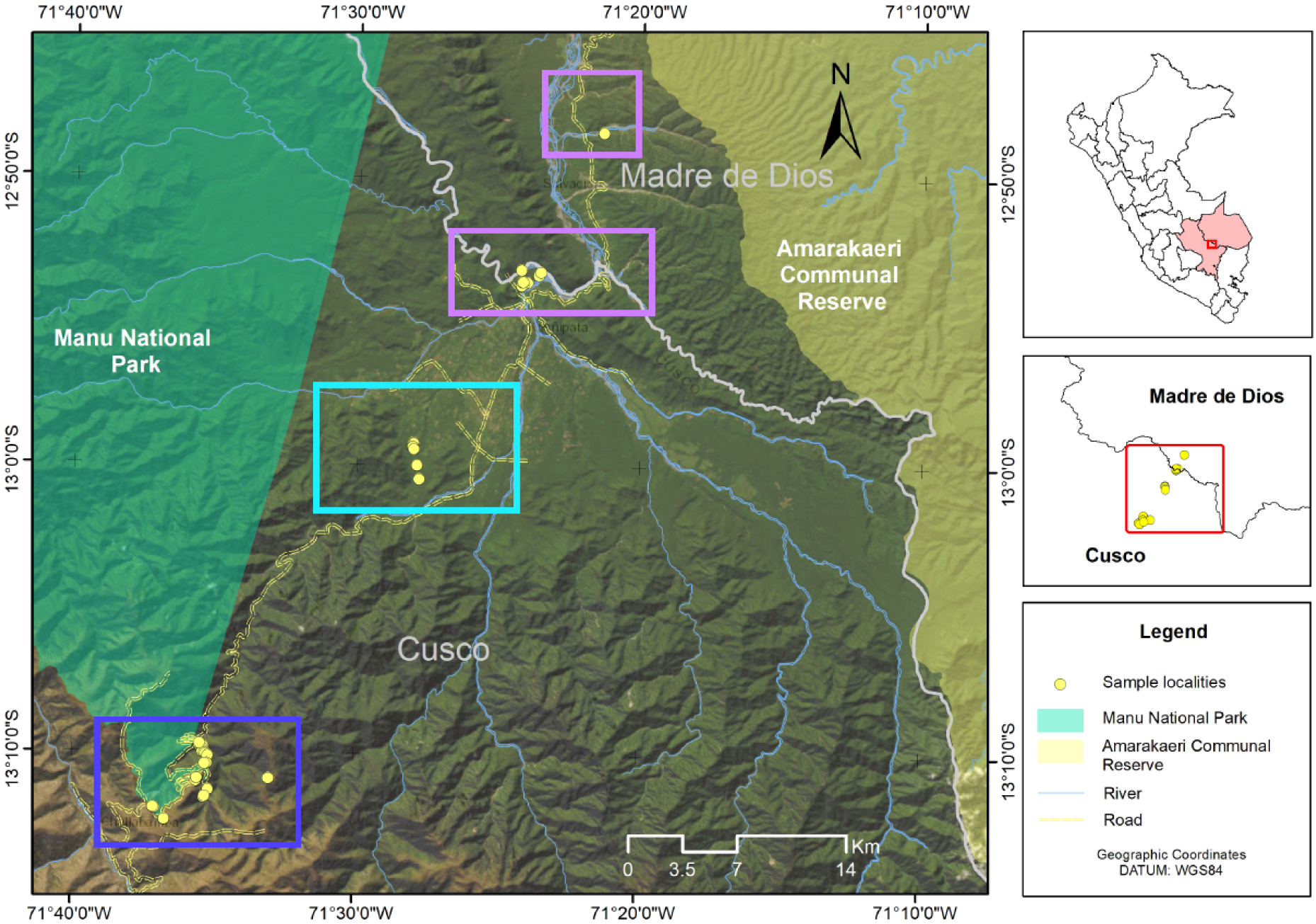
Map with the sampling sites (yellow dots) along the protected area of Madre de Dios (Cusco, Peru). The dark blue rectangle englobes the high elevation samples, the light blue rectangle englobes the intermediate elevation samples and the purple rectangles englobe the low elevation samples.

### eDNA extraction and amplification

For the environmental DNA (eDNA) extraction, we used the Qiagen DNeasy PowerSoil Pro kit following the information provided by the manufacturer. Every extracted sample was conserved at -20 °C if not immediately used. To amplify eDNA, we followed the Arcellinida-specific protocol of González-Miguéns et al. (González-Miguéns et al., 2023). We used indexed primers for subsequent demultiplexing, whose sequences are available as Supplementary Information Table S1. We purified each successfully amplified PCR product by excising the band and then quantified it using a Qubit 3 Fluorometer with dsDNA High Sensitivity (HS) assay kits (ThermoFisher). Subsequently, we standardised and pooled the samples together before sending them to the Fundación Parque Científico de Madrid for Illumina NextSeq sequencing. The demultiplexed fastq obtained can be accessed in Supplementary Information File S1.

### Data curation and tag-jumping

We followed González-Miguéns et al. (González-Miguéns et al., 2023) the pipeline for data curation, using Cutadapt v2.8 (Martin, 2011) to trim the primers and to demultiplex, then FASTQC v. 0.11.9 (Andrews, 2010) to assess the quality profiles, and finally the R package “dada2” (Callahan et al., 2016) to filter and trim the sequences. Chimeric sequences were removed and paired-ends were merged using DADA2 (Callahan et al., 2016), generating amplicon sequence variants (ASV). The information on the filtered sequences can be found in Supplementary Information Table S2. Then, we used VSEARCH v2.21.1 (Rognes et al., 2016) for the taxonomic classification, comparing our data against a curated database for eukaryotic COI (González-Miguéns et al., 2025).

After filtering the main data, we removed putative tag-jumping artefacts to avoid false positives as much as possible (Schnell et al., 2015). For that purpose, we followed the protocol suggested by ElKhouri-Vidarte et al . (2026). As well, we removed the ASVs that had an identity value for Arcellinida lower than 84%, a threshold determined empirically as explained in (González-Miguéns et al., 2024).

### Genetic analyses and clustering into OTUs

After the data curation, we aligned our sequences with those from the eukaryotic COI database (González-Miguéns et al., 2025) using MAFFT (Katoh et al., 2002) as implemented in Geneious Primer Bioinformatics Software v2.1.2023 (*Geneious | Bioinformatics Software for Sequence Data Analysis*, n.d.). We constructed a phylogenetic tree with IQ-TREE v1.6.12 (Minh et al., 2020), illustrated using FigTree v1.4.4 and R v4.3.2 (Rambaut, A., 2018).

The ASVs were clustered into OTUs using R v4.3.2 (R Core Team, 2023) with the “DECIPHER” package (Wright, 2015) using the “complete” method to obtain a proxy of species. We determined these OTUs as approximations of species, based on a barcoding gap of 0.03 as established in previous studies (ElKhouri-Vidarte et al., 2026; García-Bodelón et al., 2024; González-Miguéns et al., 2025; Useros et al., 2025). With the “ape” package (Paradis & Schliep, 2019) we transformed the ASV tree in an OTU tree.

### Drivers selection and diversity analyses

We downloaded climatic variables from Google Earth Engine (Gorelick et al., 2017) and from MapBiomas Perú (*MapBiomas*, 2026). The chosen variables were maximum and minimum temperature, actual evapotranspiration, precipitations, secondary vegetation, elevation, canopy cover and land cover. Canopy cover indicates how dense, or patchy the canopy was, while land cover is a categorical variable that reflected ecosystem types (e.g., grassland, forest etc.). To extract the data from the coordinates, we used the package “raster” (Hijmans, 2018). We assessed the correlation of the variables using the function cor() of the “stats” package (R Core Team, 2023). Because elevation, precipitation and both temperatures were highly correlated (>80%) we removed all but elevation from further analyses, based on its explanatory power established with the package “factoextra” (Alboukadel & Fabian, 2016). Elevation was divided into three sections: high elevation (ranging between 2000m and 3999m), intermediate elevation (ranging between 1000 and 1999m) and low elevation (below 1000 m) following the study from Krashevska et al. (2007) in Ecuador. To perform the ecological analyses of both alpha and beta diversity we used the following R packages: “phyloseq” (McMurdie & Holmes, 2013) as well as “ggplot2” (Wickham, 2016), “microbiome” (Lahti & Shetty, 2018), “picante” (Kembel et al., 2010), “vegan” (Oksanen et al., 2022) and “betapart” (Baselga et al., 2012). For the statistical analyses other packages as “FSA” (Ogle et al., 2023), “stats” (R Core Team, 2023) and “car” (Fox & Weisberg, 2018). The data of the drivers in each sampling site can be found in Supplementary Table S3.

## Results

From the 200 samples collected, 170 could be amplified through PCR. From those, only 47 had sequences belonging to Arcellinida, the rest consisting in sequences belonging to other microeukaryotes. In total, we filtered over 13,500,000 reads from 15,487 ASVs from those 47 samples. After removing chimeras, singletons, potential tag-jumping artefacts and non-Arcellinida sequences, we retained 2,718,374 reads from 883 ASVs that were posteriorly clustered into 113 OTUs. We obtained a median of 7.34% Arcellinida reads per sample.

While the most represented infraorder was Excentrostoma (89 OTUs and 2,140,739 reads, which represents 78.76% of the reads and 79% of the OTUs) we did also find several sequences related to Trigonopyxis (16 OTUs and 552,652 reads, 20.33% of the reads and 14% of the OTUs) and a few belonging to Hyalospheniformes (2 OTUs and 23,717 reads, 0.87% of the reads and 1.7% of the OTUs), Sphaerothecina (3 OTUs and 366 reads, 0.01% of the reads and 2.65% of the OTUs) and Cylindrothecina (3 OTUs and 450 reads, 0.02% of the reads and 2.65 % of the OTUs). High elevations had the most infraorders represented (5 infraorders, Figure 2). Both intermediate and lower elevations had three infraorders each, the latter had an almost exclusive representation of Excentrostoma, with 99.5% of the OTUs belonging to that infraorder (Figure 2).

**Figure 2:**
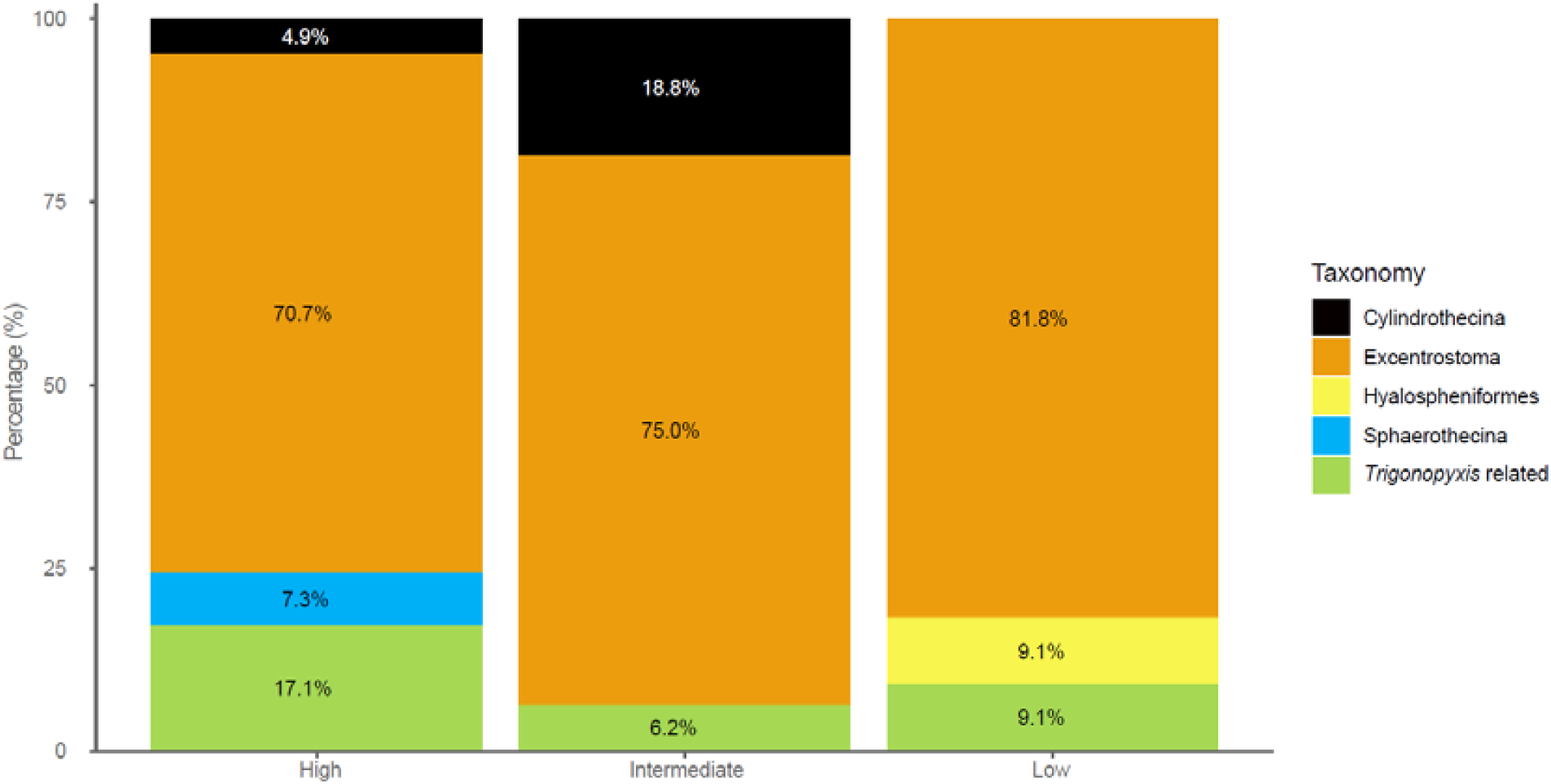
Barplot showing the proportion of each infraorder in the different sections of the elevation gradient. High elevation included samples collected above 2000 m, intermediate elevation between 1000 and 1999 m and low elevation is below 1000 m.

Along the elevation gradient we noticed significant differences in alpha diversity notably between the intermediate and the low elevations (Figure 3). Richness, phylogenetic diversi ty (PD) and Simpson’ s evenness were statistically different between those two sections (p<0.01, p<0.05 and p>0.05 respectively), richness and PD peaking at intermediate elevations while Simpson’s evenness was lower at intermediate elevations. Moreover, Simpson’s evenness was marginally lower at intermediate elevations when compared to the high elevations (p<0.08). High and low elevations did not show any statistically significant difference regarding alpha diversity (Figure 3).

**Figure 3:**
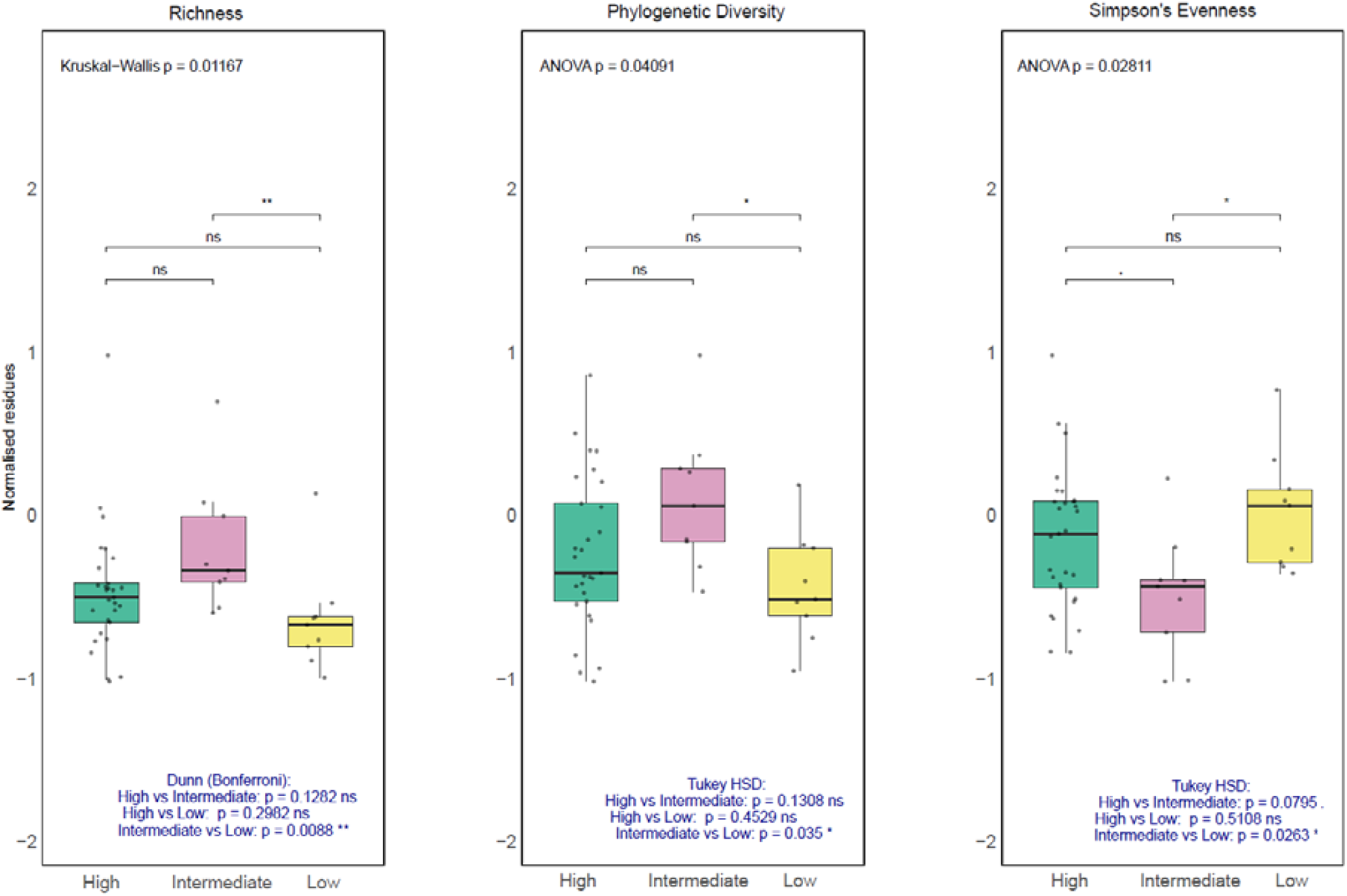
Boxplots representing different indexes of alpha diversity in the three sections of the elevation gradient. High elevation is represented in green, intermediate elevation is represented in pink and low elevation is represented in yellow. The symbol ** represents p<0. 01, * p<0.05, . 0.05<p<0. 1 and ns non-significant.

The decomposition of beta diversity showed that species turnover (i.e. species replacement) was greater and less heterogeneous at higher and lower elevations (βSIM 0.5-1, peaking at 1) than at intermediate elevations (βSIM 0-1, peaking at 0.55) (Figure 4). This result shows as well that nestedness peaks at intermediate elevations since the total beta diversity is calculated with the sum of nestedness and turnover. The differences in turnover between higher and intermediate elevations, as well as between lower and intermediate elevations were statistically significant (p<0.001, Figure 5). There was no statistically significant difference between turnover in higher and lower elevations.

**Figure 4:**
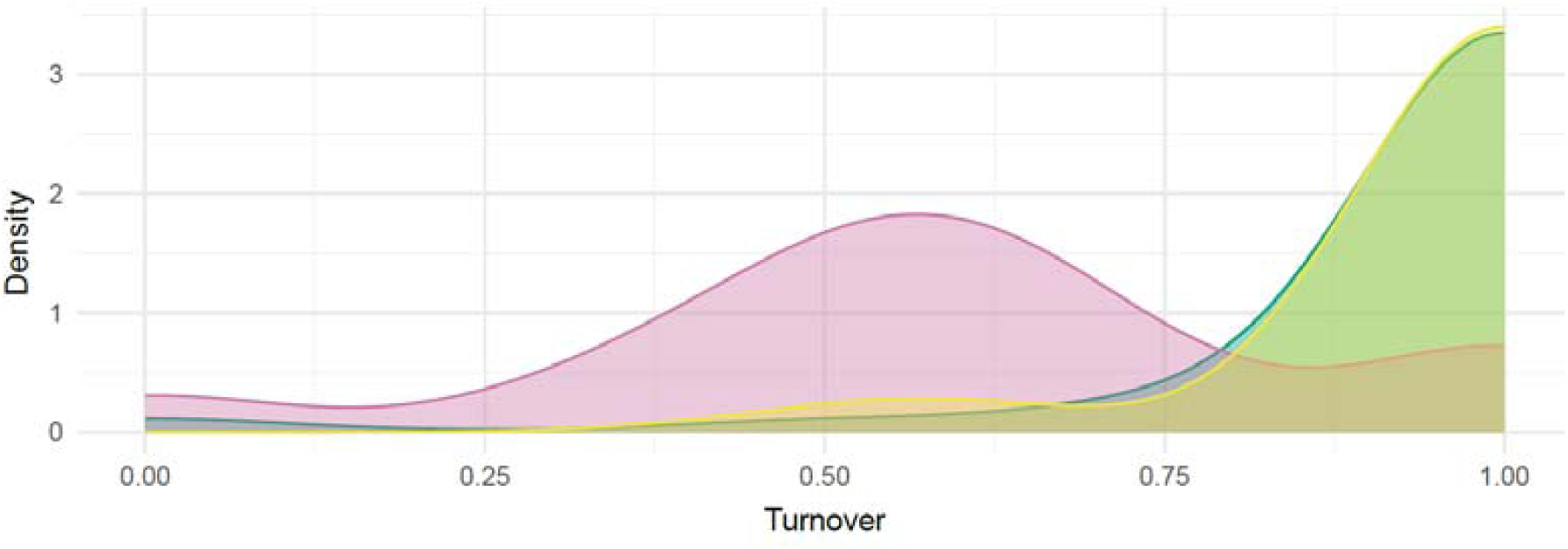
Partitioning of beta diversity showing the turnover (i. e., species replacement between communities due to environmental or spatial differences, βSIM). Kernel density curves were constructed by resampling 8 sites from each area 1000 times and computing the average βSIM. Green represents high elevations, pink represent intermediate elevations and yellow represents low elevations.

**Figure 5:**
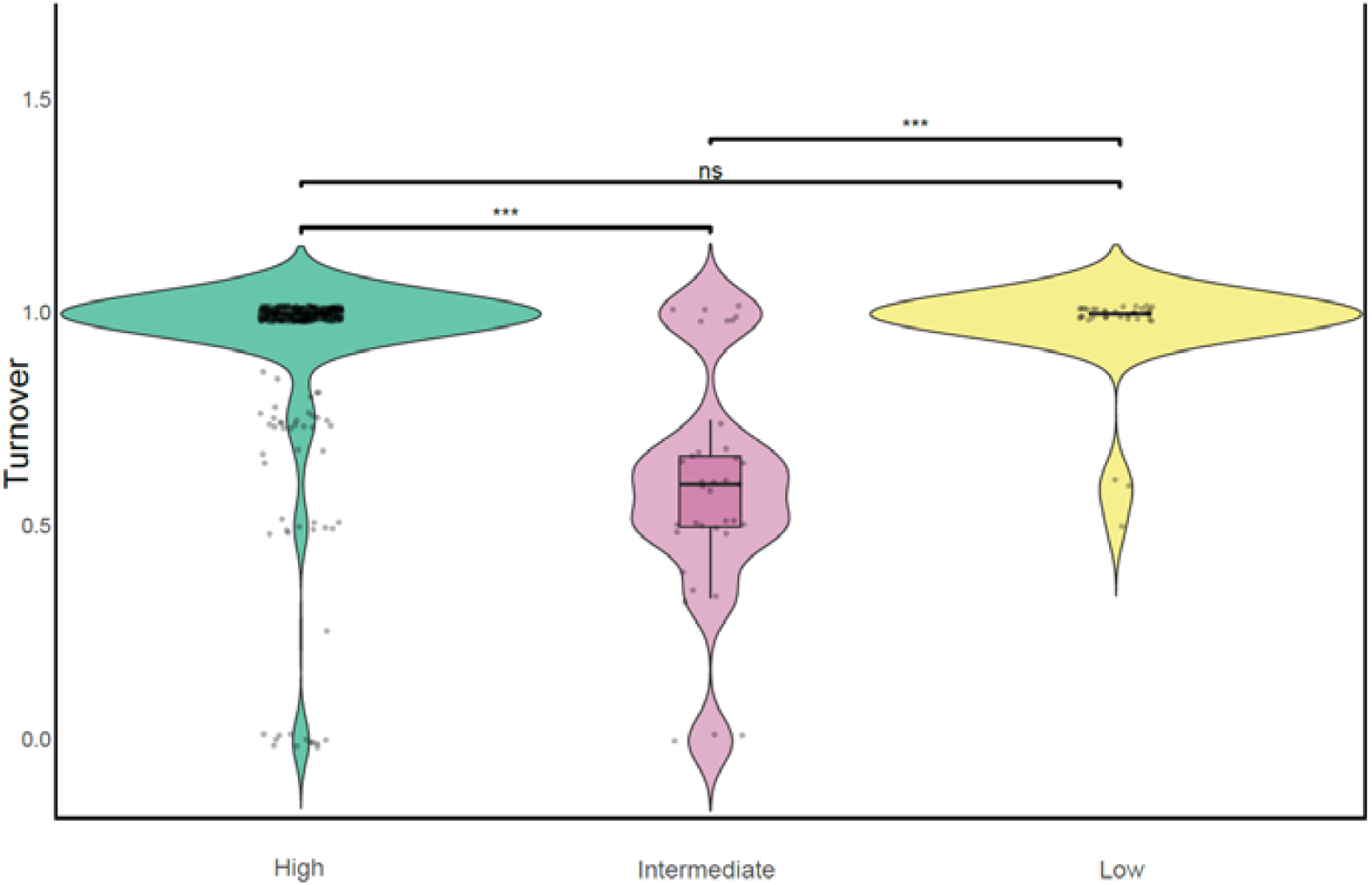
violin boxplot of the turnover in each elevation section. To compare the significance of the differences between high. Intermediate and low elevations, we applied a Kruskal-Wallis test with a Bonferroni correction. The symbol *** represents p<0. 001 and ns non-significant.

Among the variables that were tested as potential modulators of alpha and beta diversi ty, only the tree and land cover showed an effect, while other variables like temperature or humidity were never significant. Canopy cover had a significant and positive effect on PD (R2 =8.74%, p<0.05) but did not affect species richness or Simpson’s evenness (p>0.05, Table 1). Land cover was the main factor correlated with community structure (R^2^=11.63%, p<0.05) followed by canopy cover (R^2^=4.26%, p<0.01). Turnover was significantly lower at intermediate elevations (Table 1).

**Table 1:**
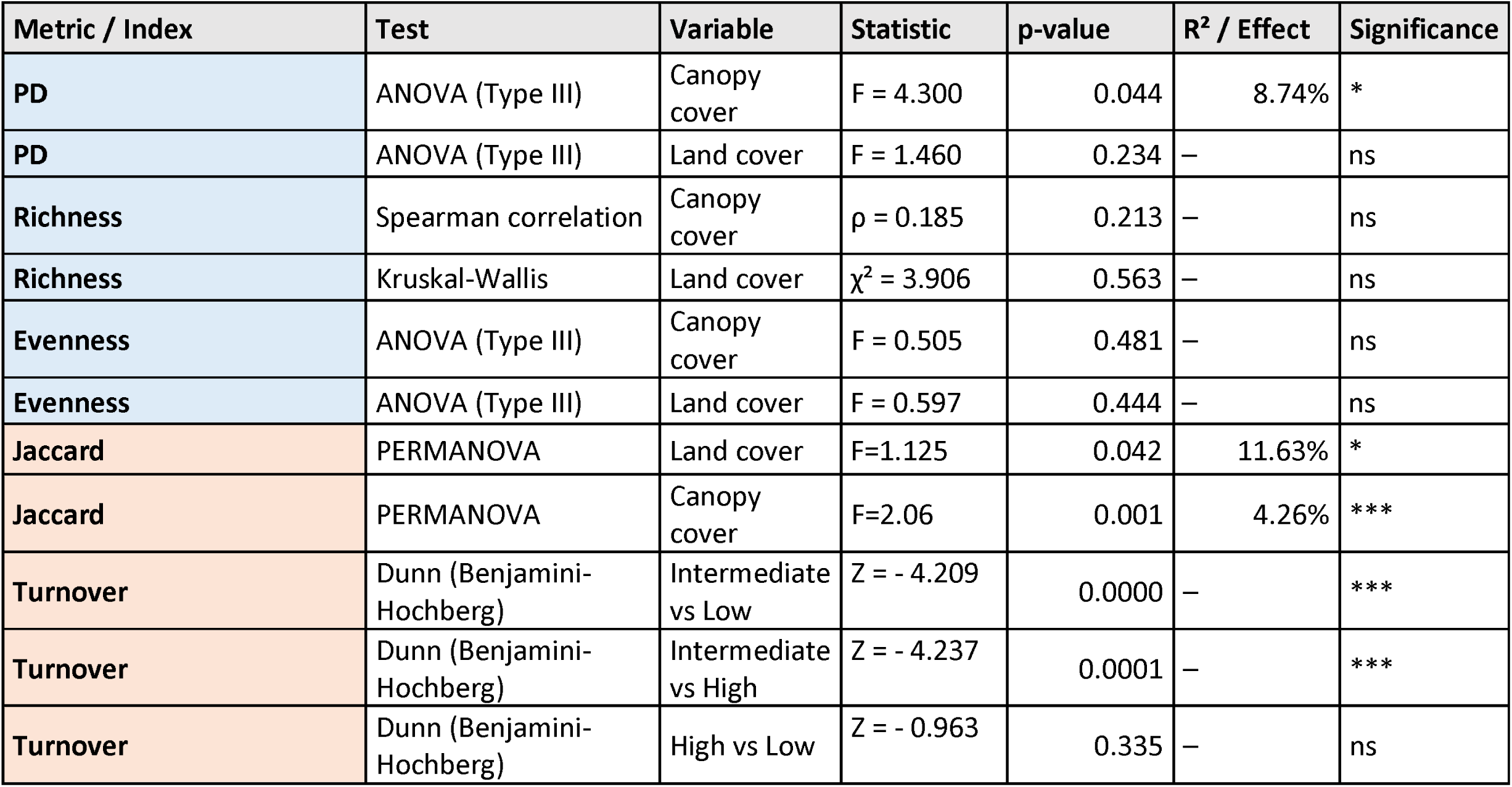
Statistic results of the effect of environmental variables in alpha (in blue) and beta diversity (in orange). The symbol *** represents p<0. 001, * p<0.05, . 0. 05<p<0. 1 and ns non-significant. For turnover, values of the effect represent mean differences between habitat pairs (Games-Howell test). The test was chosen depending on the distribution of the data and the nature of the explanatory variables (i. e., categorical vs continuous).

## Discussion

### General aspects of Arcellinida diversity in the Neotropics

In this study we examined soil Arcellinida communities in one of the world’s diversity hotspots, assessing the changes in the communities through environmental factors. To our knowledge, this is the first eDNA survey on Arcellinida conducted in the Andean mountain range and in any Neotropical forest. However, the density of Arcellinida was surprisingly lower than expected (7.34% of Arcellinida reads out of the total reads during sequencing) since we did not obtain any Arcellinida sequence in the majority of the samples (only in 47 out of 170 sequenced samples). This contrasts also with the findings of Kang and Dumack (2026) in temperate forests and found local dominance of Arcellinida, even though they did not use specific primers, and with the findings of González-Miguéns et al. (2024) in extratropical environments using the same primer set. They proposed that Arcellinida could be the most abundant eukaryotic taxa in certain samples based on an approximation built on environmental RNA.

Diversity encountered was also surprisingly low. Indeed, a comparable study in Mediterranean woodlands retrieved almost 700 OTUs within 120 samples, while in this study we found only 113 OTUs in 200 samples. This pattern of low number of species (113 OTUs) but higher diversity (5 infraorders represented in our samples) is consistent with the findings of Krashevska et al. (2007) in the Ecuatorian Neotropics. While this low diversity certainly corresponds with a biological reality, we are aware that our primer set biases diversity against infraorder Hyalospheniformes (González Miguéns et al., 2024), a group that is present in the Neotropics and has been detected in Neotropical rainforests using morphological identification (Alcino et al., 2026; Krashevska et al., 2007). In that sense, it can also be possible that the primer set used here presents mismatches with other species present in Neotropical areas. In any case, the low diversity encountered in this study does not support the hypothesis that Neotropical rainforests are hotspots for Arcellinida diversity, and suggests rather the opposite. Although the latitudinal diversity gradient is widely observed across many taxa, it is not universal, and substantial variation occurs among taxonomic groups, habitats and spatial scales (Willig et al., 2003). A recent study that recompiled data from pro- and eukaryotic microbes, including soil fauna, showed no clear trend towards the tropics (Gao et al., 2026). Whether the low Arcellinida diversity found in the Kosñipata Valley can be extrapolated to all the wet tropics remains to be tested. Still, our results show that Arcellinida deviate from a pattern that is generally accepted for plants and animals.

As it usually happens in soil-based studies, the majority of the OTUs recovered belonged to the infraorder Excentrostoma (ElKhouri-Vidarte et al., 2026; González-Miguéns et al., 2023). But other infraorders such as Hyalospheniformes or sequences related to *Trigonopyxis* were still relatively abundant while Sphaerothecina and Cylindrothecina were present in smaller numbers. The *Trigonopyxis* group has often been reported in outside of the temperate areas, but seems to be more represented in Neotropical ecosystems (Bobrov et al., 2013; Schulz et al., 2018). While this overall composition broadly resembl es that of extratropical forests (ElKhouri-Vidarte et al., 2026), the notable prominence of sequences related to *Trigonopyxis* points to a distinct biogeographic signal characteristic of Neotropical ecosystems.

### Patterns of diversity along the elevation gradient

Arcellinida diversity in the Andean mountain range differed between elevation sections. The intermediate elevations tended to be more diverse, although there was only a statistically significant difference between the intermediate and the low elevations. For richness and PD there was a unimodal distribution along the elevation sections. This is consistent with findings in temperate and Mediterranean climates, where Arcellinida richness peaked at intermediate elevations with a milder and more humid climate (ElKhouri-Vidarte et al., 2025; Lamentowicz et al., 2013). Under tropical climates devoid of dry season, results can be similar (Krashevska et al., 2007) but the mechanisms behind may be different and need to be explored.

When decomposing the beta diversity into nestedness and turnover (Baselga et al., 2012), drastic differences appear between intermediate and extreme (high or low) elevations. Turnover was significantly higher at extreme (higher and lower) elevations than at intermediate ones, and, importantly, intermediate elevations showed a broader range of turnover values. This pattern suggests that communities at intermediate elevations are not simply transitional between Andean and Amazonian assemblages. Instead, they may reflect the existence of multiple local environmental conditions supporting distinct community configurations. Indeed, some studies have found that high nestedness dominates in unstable regions that have endured processes of extinction and subsequent colonisation (Zhang et al., 2022). Intermediate elevations are also more heterogeneous (lower evenness), creating a nested structure in which impoverished, disturbed communities are subsets of richer, undisturbed communities (Ulrich et al., 2009). In Arcellinida, such dynamics have been shown in peat bogs (Koenig et al., 2017) where disturbances were experimentally induced droughts. The occurrence of flash floods and/or landslides is pervasive in the Kosñipata Valley, especially around 1700 m where a river knickpoint causes geological instability, with verified consequences on biodiversity (Clark et al., 2016).

Contrarily, higher elevations host more specialised species that can endure strong environmental filters in adequate microniches, which promotes higher turnover (Jiang et al., 2023, Wang et al., 2012). In turn, at lower elevations the productivity is higher because of the high humidity and temperatures, and there is a higher niche specialisation promoted by a strong competition that enhances a high turnover (Zhang et al., 2024).

### Above ground effects on below-ground diversity

At all elevations, Arcellinida distribution is mediated by the canopy cover, as it affects both the PD and beta diversity (i.e., species composition, R^2^=8.74% and R^2^=4.26% respectively). Canopy cover is indirectly related to ecological filter since it influences through changes in the litter, soil moisture, pH, and even oxygen, sugars and organic acids due to the roots and its exudates (Berg et al., 2015; Okubo et al., 2016; Rosier et al., 2015). An important canopy cover creates buffered microclimatic conditions (Verdonck et al., 2025), promoting environmental stability and potentially favouring the persistence of a wider range of evolutionary lineages. Indeed, testate amoebae have been suggested to be more diverse and abundant in soils with high moisture, organic content and slow decomposition rates (Krashevska et al., 2009).

At a broader landscape scale, land cover was the strongest predictor of Arcellinida community composition (R^2^= 11.63%) which is consistent with other studies conducted in extratropical freshwater ecosystems (Soler-Zamora et al., 2026). The different land-cover categories in our dataset, including forests, grasslands, agricultural mosaics and non-forested areas, likely represent distinct environmental filters that shape community composition. These habitats differ not only in vegetation structure, but also in litter production, moisture retention, soil properties and resource availability.

As well, it is consistent with previous studies that have found vegetation as one of the main factors that modulates Arcellinida diversity (Tsyganov et al., 2022). These drivers have been demonstrated to affect testate amoebae communities (Krashevska et al., 2017) and canopy cover may create buffered microclimatic conditions (Verdonck et al., 2025), promoting environmental stability and potentially favouring the persistence of a wider range of evolutionary lineages.

## Conclusions

This study shows the unexpected low diversity of Arcellinida in a world diversity hotspot. We found that Neotropical communities differed in their composition from extratropical forest litter, as we found a high representation of species related to *Trigonopyxis* and some taxa from the Hyalospheniformes clade. While our results show the effects of elevation on diversity, vegetation, expressed in terms of land and canopy cover, seem to play the greatest role in explaining Arcellinida diversity in the Kosñipata Valley. Nevertheless, as opposed to Mediterranean ecosystems, biotic interactions seem to drive biodiversity rather than direct influence of climate. Our findings also reiterate the importance of vegetation in modulating Arcellinida communities and diversity.

Further research is needed in different ecosystems to improve our understanding of soil litter and its influence on testate amoebae. This should include the contribution of canopy cover and an integrated approach to above- and belowground processes within one ecosystem.

Future work across a broader range of tropical ecosystems will be essential to determine whether this unexpectedly low Arcellinida diversity is a localized phenomenon or a widespread feature of the wet tropics.

## Acknowledgments

We acknowledge the grant IND2022/AMB-23616 awarded by the community of Madrid in the program “doctorados industriales” as well as a grant from the Spanish Government “Generación de Conocimiento” PID2021-128499NB-I00 10.13039/501100011033/, (MCIU/AEI/ FEDER, UE).

## Supplementary Information

File S1: Illumina paired-end sequences in fastq format, obtained after the demultiplexing.

**Table S1:**
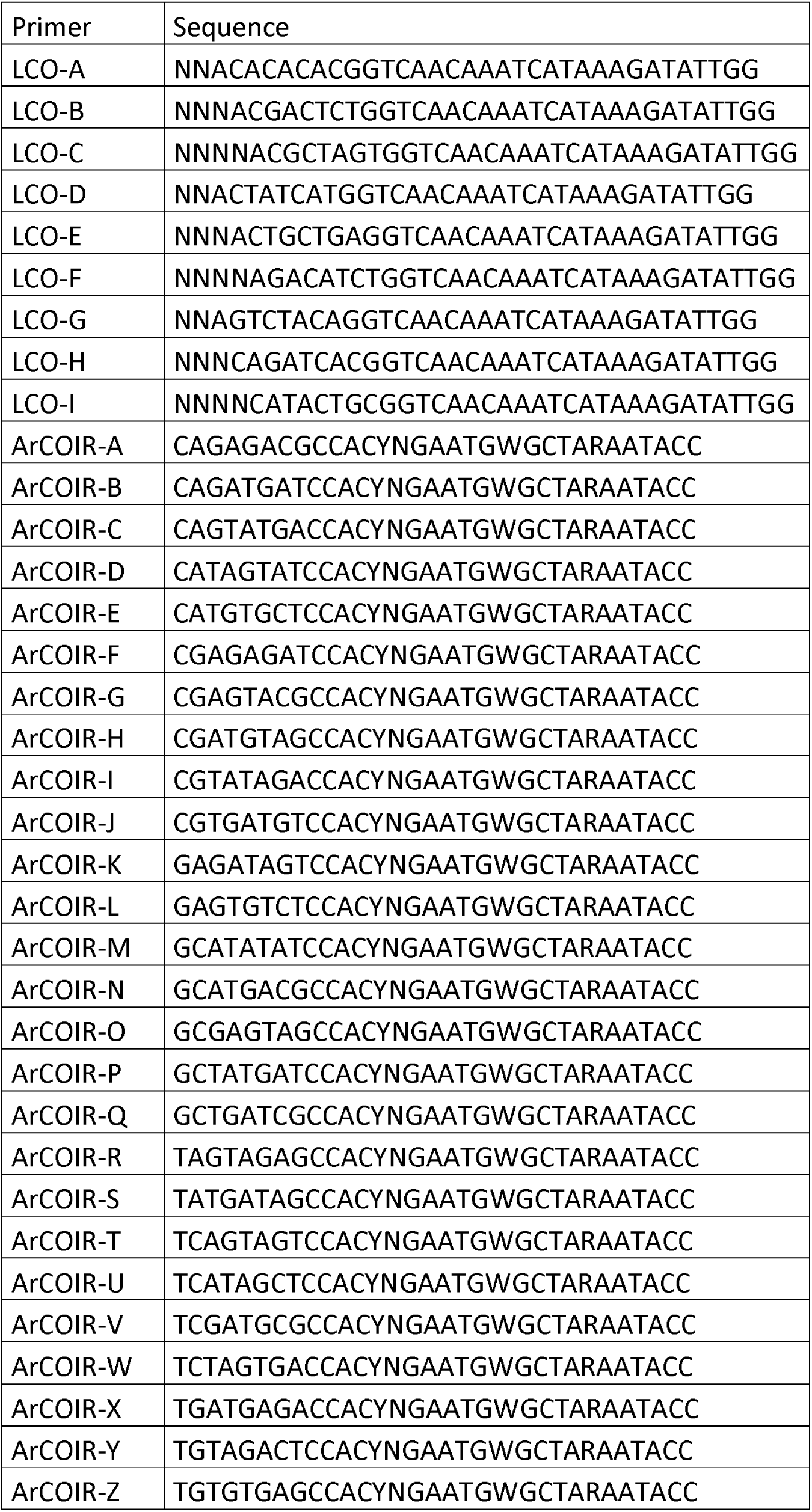
Indexed primers used in this study and their sequence. Excluding the initial Ns, the eight first nucleotides correspond to the index, and the rest constitute the primer sequence.

**Table S2:** Information of the filtered sequences after DADA2 cleaning step.

| Sample | input | filtered | denoisedF | denoisedR | merged | nonchim | perc |
| --- | --- | --- | --- | --- | --- | --- | --- |
| LCO-A-ArCOIR-A_Am001 | 345916 | 335528 | 334836 | 334517 | 2916 | 2746 | 0.79 |
| LCO-A-ArCOIR-B_Am002 | 1203 | 951 | 823 | 846 | 68 | 68 | 5.65 |
| LCO-A-ArCOIR-C_Am003 | 1258 | 1033 | 909 | 958 | 18 | 18 | 1.43 |
| LCO-A-ArCOIR-D_Am004 | 481876 | 477644 | 475525 | 476954 | 588 | 580 | 0.12 |
| LCO-A-ArCOIR-E_Am005 | 557649 | 548024 | 545853 | 543435 | 963 | 915 | 0.16 |
| LCO-A-ArCOIR-F_Am006 | 337811 | 313826 | 312000 | 312604 | 14530 | 7296 | 2.16 |
| LCO-A-ArCOIR-G_Am007 | 657042 | 636592 | 634261 | 634738 | 29074 | 23327 | 3.55 |
| LCO-A-ArCOIR-H_Am008 | 405009 | 400643 | 397533 | 398442 | 1052 | 978 | 0.24 |
| LCO-A-ArCOIR-I_Am009 | 269 | 198 | 142 | 176 | 48 | 48 | 17.84 |
| LCO-A-ArCOIR-J_Am010 | 541237 | 521933 | 520751 | 518938 | 1132 | 1113 | 0.21 |
| LCO-A-ArCOIR-K_Am011 | 160 | 141 | 104 | 132 | 0 | 0 | 0 |
| LCO-A-ArCOIR-L_Am012 | 741 | 738 | 703 | 716 | 4 | 4 | 0.54 |
| LCO-A-ArCOIR-M_Am013 | 19 | 19 | 18 | 18 | 0 | 0 | 0 |
| LCO-A-ArCOIR-N_Am014 | 273665 | 238951 | 238321 | 238300 | 62709 | 43760 | 15.99 |
| LCO-A-ArCOIR-O_Am015 | 29875 | 28927 | 28845 | 28713 | 1145 | 905 | 3.03 |
| LCO-A-ArCOIR-P_Am016 | 704 | 672 | 591 | 624 | 4 | 4 | 0.57 |
| LCO-A-ArCOIR-Q_Am017 | 125 | 121 | 64 | 81 | 0 | 0 | 0 |
| LCO-A-ArCOIR-R_Am018 | 212095 | 207921 | 207424 | 207667 | 2220 | 2214 | 1.04 |
| LCO-A-ArCOIR-S_Am019 | 7 | 7 | 5 | 5 | 0 | 0 | 0 |
| LCO-A-ArCOIR-T_Am020 | 9178 | 8805 | 8727 | 8717 | 234 | 231 | 2.52 |
| LCO-A-ArCOIR-U_Am021 | 178897 | 175345 | 175188 | 174830 | 1743 | 1742 | 0.97 |
| LCO-A-ArCOIR-V_Am022 | 247 | 194 | 147 | 148 | 79 | 75 | 30.36 |
| LCO-A-ArCOIR-W_Am023 | 4 | 4 | 3 | 3 | 0 | 0 | 0 |
| LCO-A-ArCOIR-X_Am024 | 0 | 0 | 310663 | 310811 | 4046 | 3972 | Inf |
| LCO-A-ArCOIR-Y_Am025 | 0 | 0 | 422 | 423 | 156 | 156 | Inf |
| LCO-A-ArCOIR-Z_Am026 | 318184 | 311181 | 203 | 199 | 0 | 0 | 0 |
| LCO-B-ArCOIR-A_Am027 | 616 | 488 | 290 | 294 | 0 | 0 | 0 |
| LCO-B-ArCOIR-B_Am028 | 268 | 231 | 193485 | 193919 | 26 | 26 | 9.7 |
| LCO-B-ArCOIR-C_Am029 | 358 | 337 | 587 | 551 | 12 | 12 | 3.35 |
| LCO-B-ArCOIR-D_Am030 | 196427 | 194049 | 1965 | 1992 | 833 | 754 | 0.38 |
| LCO-B-ArCOIR-E_Am031 | 720 | 667 | 965 | 975 | 127 | 127 | 17.64 |
| LCO-B-ArCOIR-F_Am032 | 13443 | 2216 | 982 | 993 | 93 | 51 | 0.38 |
| LCO-B-ArCOIR-G_Am033 | 1370 | 1099 | 855 | 876 | 9 | 9 | 0.66 |
| LCO-B-ArCOIR-H_Am034 | 1160 | 1078 | 91 | 80 | 0 | 0 | 0 |
| LCO-B-ArCOIR-I_Am035 | 920 | 894 | 15 | 20 | 0 | 0 | 0 |
| LCO-B-ArCOIR-J_Am036 | 116 | 111 | 186636 | 186023 | 21 | 21 | 18.1 |
| LCO-B-ArCOIR-K_Am037 | 35 | 29 | 129 | 129 | 0 | 0 | 0 |
| LCO-B-ArCOIR-L_Am038 | 188673 | 187489 | 59 | 70 | 29 | 29 | 0.02 |
| LCO-B-ArCOIR-M_Am039 | 132 | 131 | 11 | 9 | 5 | 5 | 3.79 |
| LCO-B-ArCOIR-N_Am040 | 98 | 84 | 242 | 243 | 2 | 2 | 2.04 |
| LCO-B-ArCOIR-O_Am041 | 13 | 13 | 11 | 14 | 6 | 6 | 46.15 |
| LCO-B-ArCOIR-P_Am042 | 277 | 267 | 35 | 37 | 0 | 0 | 0 |
| LCO-B-ArCOIR-Q_Am043 | 24 | 23 | 12 | 15 | 0 | 0 | 0 |
| LCO-B-ArCOIR-R_Am044 | 50 | 48 | 146 | 142 | 0 | 0 | 0 |
| LCO-B-ArCOIR-S_Am045 | 0 | 0 | 49 | 54 | 4 | 4 | Inf |
| LCO-B-ArCOIR-T_Am046 | 18 | 16 | 41 | 44 | 0 | 0 | 0 |
| LCO-B-ArCOIR-U_Am047 | 165 | 165 | 223689 | 223831 | 642 | 642 | 389.09 |
| LCO-B-ArCOIR-V_Am048 | 71 | 63 | 899 | 931 | 17 | 16 | 22.54 |
| LCO-B-ArCOIR-W_Am049 | 0 | 0 | 721 | 744 | 7 | 7 | Inf |
| LCO-B-ArCOIR-X_Am050 | 0 | 0 | 1253 | 1267 | 12 | 12 | Inf |
| LCO-B-ArCOIR-Y_Am051 | 0 | 0 | 254281 | 254601 | 1707 | 1707 | Inf |
| LCO-B-ArCOIR-Z_Am052 | 68 | 53 | 2930 | 2858 | 1383 | 1284 | 1888.24 |
| LCO-C-ArCOIR-A_Am053 | 226321 | 224621 | 257538 | 257366 | 3447 | 2553 | 1.13 |
| LCO-C-ArCOIR-B_Am054 | 1374 | 993 | 541798 | 542245 | 793 | 774 | 56.33 |
| LCO-C-ArCOIR-C_Am055 | 849 | 807 | 51 | 58 | 15 | 15 | 1.77 |
| LCO-C-ArCOIR-D_Am056 | 1392 | 1355 | 213909 | 213886 | 1083 | 995 | 71.48 |
| LCO-C-ArCOIR-E_Am057 | 257607 | 254897 | 392936 | 393223 | 671 | 671 | 0.26 |
| LCO-C-ArCOIR-F_Am058 | 12807 | 3158 | 92160 | 92679 | 131 | 131 | 1.02 |
| LCO-C-ArCOIR-G_Am059 | 268077 | 258523 | 60 | 63 | 0 | 0 | 0 |
| LCO-C-ArCOIR-H_Am060 | 550144 | 543567 | 136 | 137 | 24 | 24 | 0 |
| LCO-C-ArCOIR-I_Am061 | 103 | 73 | 232 | 237 | 0 | 0 | 0 |
| LCO-C-ArCOIR-J_Am062 | 217010 | 214175 | 355964 | 354676 | 1040 | 1036 | 0.48 |
| LCO-C-ArCOIR-K_Am063 | 396884 | 393614 | 59 | 49 | 0 | 0 | 0 |
| LCO-C-ArCOIR-L_Am064 | 93836 | 92702 | 301499 | 298993 | 3 | 3 | 0 |
| LCO-C-ArCOIR-M_Am065 | 73 | 70 | 3 | 1 | 0 | 0 | 0 |
| LCO-C-ArCOIR-N_Am066 | 207 | 174 | 50 | 62 | 0 | 0 | 0 |
| LCO-C-ArCOIR-O_Am067 | 261 | 255 | 476979 | 475115 | 82 | 82 | 31.42 |
| LCO-C-ArCOIR-P_Am068 | 358736 | 356134 | 83 | 87 | 5 | 5 | 0 |
| LCO-C-ArCOIR-Q_Am069 | 92 | 80 | 7 | 7 | 0 | 0 | 0 |
| LCO-C-ArCOIR-R_Am070 | 304320 | 302508 | 124 | 134 | 0 | 0 | 0 |
| LCO-C-ArCOIR-S_Am071 | 6 | 6 | 455763 | 456423 | 34749 | 20421 | 340350 |
| LCO-C-ArCOIR-T_Am072 | 76 | 75 | 466 | 478 | 30 | 14 | 18.42 |
| LCO-C-ArCOIR-U_Am073 | 479743 | 477694 | 345968 | 346489 | 584 | 575 | 0.12 |
| LCO-C-ArCOIR-V_Am074 | 138 | 120 | 177574 | 178011 | 5034 | 3310 | 2398.55 |
| LCO-C-ArCOIR-W_Am075 | 7 | 7 | 155448 | 155145 | 912 | 912 | 13028.57 |
| LCO-C-ArCOIR-X_Am076 | 0 | 0 | 4696 | 4559 | 1546 | 1379 | Inf |
| LCO-C-ArCOIR-Y_Am077 | 2 | 0 | 670577 | 668040 | 82270 | 25574 | 1278700 |
| LCO-C-ArCOIR-Z_Am078 | 177 | 157 | 514327 | 515003 | 68577 | 20664 | 11674.58 |
| LCO-D-ArCOIR-A_Am079 | 475356 | 458866 | 196 | 226 | 97 | 89 | 0.02 |
| LCO-D-ArCOIR-B_Am080 | 760 | 544 | 501 | 505 | 0 | 0 | 0 |
| LCO-D-ArCOIR-C_Am081 | 352762 | 347177 | 87 | 98 | 31 | 31 | 0.01 |
| LCO-D-ArCOIR-D_Am082 | 180388 | 178422 | 70 | 80 | 0 | 0 | 0 |
| LCO-D-ArCOIR-E_Am083 | 174972 | 156423 | 137 | 137 | 0 | 0 | 0 |
| LCO-D-ArCOIR-F_Am084 | 16627 | 5015 | 85189 | 85007 | 3520 | 3453 | 20.77 |
| LCO-D-ArCOIR-G_Am085 | 714027 | 672284 | 715 | 708 | 125 | 120 | 0.02 |
| LCO-D-ArCOIR-H_Am086 | 540603 | 516490 | 288 | 309 | 0 | 0 | 0 |
| LCO-D-ArCOIR-I_Am087 | 261 | 240 | 71 | 66 | 0 | 0 | 0 |
| LCO-D-ArCOIR-J_Am088 | 565 | 543 | 109 | 121 | 3 | 3 | 0.53 |
| LCO-D-ArCOIR-K_Am089 | 121 | 105 | 34 | 48 | 3 | 3 | 2.48 |
| LCO-D-ArCOIR-L_Am090 | 88 | 87 | 157 | 165 | 11 | 11 | 12.5 |
| LCO-D-ArCOIR-M_Am091 | 142 | 141 | 66368 | 66315 | 22518 | 7253 | 5107.75 |
| LCO-D-ArCOIR-N_Am092 | 87871 | 85952 | 126 | 133 | 0 | 0 | 0 |
| LCO-D-ArCOIR-O_Am093 | 790 | 741 | 84 | 92 | 3 | 3 | 0.38 |
| LCO-D-ArCOIR-P_Am094 | 365 | 351 | 131 | 138 | 4 | 4 | 1.1 |
| LCO-D-ArCOIR-Q_Am095 | 116 | 92 | 162 | 185 | 10 | 10 | 8.62 |
| LCO-D-ArCOIR-R_Am096 | 147 | 139 | 109652 | 109356 | 1146 | 1137 | 773.47 |
| LCO-D-ArCOIR-S_Am097 | 0 | 0 | 162014 | 161908 | 455 | 454 | Inf |
| LCO-D-ArCOIR-T_Am098 | 60 | 59 | 6750 | 6793 | 2680 | 1888 | 3146.67 |
| LCO-D-ArCOIR-U_Am099 | 207 | 193 | 606 | 617 | 48 | 48 | 23.19 |
| LCO-D-ArCOIR-V_Am100 | 71378 | 66795 | 474 | 431 | 22 | 9 | 0.01 |
| LCO-D-ArCOIR-W_Am101 | 0 | 0 | 20 | 21 | 0 | 0 | NA |
| LCO-D-ArCOIR-X_Am102 | 0 | 0 | 42 | 48 | 0 | 0 | NA |
| LCO-D-ArCOIR-Y_Am103 | 0 | 0 | 3 | 11 | 0 | 0 | NA |
| LCO-D-ArCOIR-Z_Am104 | 159 | 150 | 8 | 9 | 0 | 0 | 0 |
| LCO-E-ArCOIR-A_Am105 | 131 | 123 | 93 | 83 | 2 | 2 | 1.53 |
| LCO-E-ArCOIR-B_Am106 | 190 | 162 | 17 | 27 | 13 | 13 | 6.84 |
| LCO-E-ArCOIR-C_M0001 | 235 | 220 | 49 | 54 | 0 | 0 | 0 |
| LCO-E-ArCOIR-D_M0006 | 112534 | 110027 | 114399 | 113954 | 70 | 70 | 0.06 |
| LCO-E-ArCOIR-E_M0011 | 166578 | 162482 | 18 | 32 | 0 | 0 | 0 |
| LCO-E-ArCOIR-F_M0016 | 10855 | 7031 | 5 | 12 | 0 | 0 | 0 |
| LCO-E-ArCOIR-G_M0020 | 831 | 728 | 41 | 52 | 0 | 0 | 0 |
| LCO-E-ArCOIR-H_M0023 | 564 | 538 | 15 | 14 | 6 | 6 | 1.06 |
| LCO-E-ArCOIR-I_M0030 | 31 | 28 | 4 | 4 | 0 | 0 | 0 |
| LCO-E-ArCOIR-J_M0032 | 80 | 73 | 15 | 23 | 0 | 0 | 0 |
| LCO-E-ArCOIR-K_M0036 | 25 | 22 | 102 | 108 | 10 | 3 | 12 |
| LCO-E-ArCOIR-L_M0038 | 16 | 16 | 317 | 323 | 95 | 92 | 575 |
| LCO-E-ArCOIR-M_M0048 | 103 | 102 | 38 | 53 | 2 | 2 | 1.94 |
| LCO-E-ArCOIR-N_M0049 | 42 | 37 | 147 | 152 | 6 | 6 | 14.29 |
| LCO-E-ArCOIR-O_M0050 | 1 | 0 | 324 | 339 | 11 | 11 | 1100 |
| LCO-E-ArCOIR-P_M0051 | 79 | 75 | 278 | 259 | 69 | 69 | 87.34 |
| LCO-E-ArCOIR-Q_M0053 | 115626 | 114824 | 268304 | 268443 | 1548 | 1485 | 1.28 |
| LCO-E-ArCOIR-R_M0055 | 37 | 36 | 230288 | 229773 | 15727 | 5388 | 14562.16 |
| LCO-E-ArCOIR-S_M0057 | 0 | 0 | 1102 | 1097 | 7 | 7 | Inf |
| LCO-E-ArCOIR-T_M0064 | 15 | 15 | 44 | 47 | 0 | 0 | 0 |
| LCO-E-ArCOIR-U_M0065 | 62 | 61 | 4 | 5 | 0 | 0 | 0 |
| LCO-E-ArCOIR-V_M0070 | 31 | 31 | 7 | 10 | 0 | 0 | 0 |
| LCO-E-ArCOIR-W_M0075 | 0 | 0 | 47 | 64 | 17 | 17 | Inf |
| LCO-E-ArCOIR-X_M0076 | 5 | 5 | 80 | 71 | 0 | 0 | 0 |
| LCO-E-ArCOIR-Y_M0077 | 0 | 0 | 352929 | 352503 | 529 | 524 | Inf |
| LCO-E-ArCOIR-Z_M0078 | 33 | 31 | 61786 | 61804 | 540 | 534 | 1618.18 |
| LCO-F-ArCOIR-A_M0080 | 150 | 136 | 18 | 18 | 0 | 0 | 0 |
| LCO-F-ArCOIR-B_M0082 | 901 | 366 | 4 | 7 | 0 | 0 | 0 |
| LCO-F-ArCOIR-C_M0083 | 85 | 73 | 8 | 12 | 0 | 0 | 0 |
| LCO-F-ArCOIR-D_M0084 | 188 | 180 | 27 | 27 | 18 | 18 | 9.57 |
| LCO-F-ArCOIR-E_M0085 | 421 | 391 | 54 | 62 | 6 | 6 | 1.43 |
| LCO-F-ArCOIR-F_M0086 | 509 | 313 | 7 | 11 | 0 | 0 | 0 |
| LCO-F-ArCOIR-G_M0087 | 273155 | 270251 | 150 | 163 | 0 | 0 | 0 |
| LCO-F-ArCOIR-H_M0088 | 238209 | 231029 | 1243 | 1273 | 622 | 562 | 0.24 |
| LCO-F-ArCOIR-I_M0091 | 1141 | 1121 | 162 | 190 | 0 | 0 | 0 |
| LCO-F-ArCOIR-J_M0092 | 68 | 63 | 358 | 342 | 29 | 29 | 42.65 |
| LCO-F-ArCOIR-K_M0093 | 9 | 9 | 418 | 397 | 22 | 22 | 244.44 |
| LCO-F-ArCOIR-L_M0097 | 18 | 16 | 15339 | 15235 | 2095 | 1825 | 10138.89 |
| LCO-F-ArCOIR-M_M0098 | 0 | 0 | 2403 | 2396 | 1119 | 901 | Inf |
| LCO-F-ArCOIR-N_M0099 | 82 | 79 | 2551 | 2595 | 1224 | 933 | 1137.8 |
| LCO-F-ArCOIR-O_M0101 | 105 | 104 | 12951 | 13119 | 5894 | 4319 | 4113.33 |
| LCO-F-ArCOIR-P_M0102 | 358276 | 355054 | 58 | 55 | 0 | 0 | 0 |
| LCO-F-ArCOIR-Q_M0104 | 63277 | 62307 | 21 | 22 | 0 | 0 | 0 |
| LCO-F-ArCOIR-R_M0106 | 38 | 26 | 2 | 8 | 0 | 0 | 0 |
| LCO-F-ArCOIR-S_M106B | 0 | 0 | 1 | 1 | 0 | 0 | NA |
| LCO-F-ArCOIR-T_M106S | 9 | 8 | 74 | 83 | 5 | 5 | 55.56 |
| LCO-F-ArCOIR-U_M0108 | 24 | 24 | 1 | 1 | 0 | 0 | 0 |
| LCO-F-ArCOIR-V_M0109 | 38 | 35 | 68 | 67 | 2 | 2 | 5.26 |
| LCO-F-ArCOIR-W_M0110 | 0 | 0 | 15 | 16 | 0 | 0 | NA |
| LCO-F-ArCOIR-X_M0114 | 0 | 0 | 19 | 22 | 0 | 0 | NA |
| LCO-F-ArCOIR-Y_M0115 | 116 | 74 | 171163 | 172105 | 151 | 147 | 126.72 |
| LCO-F-ArCOIR-Z_M0116 | 22 | 20 | 33 | 38 | 4 | 4 | 18.18 |
| LCO-G-ArCOIR-A_M0118 | 225 | 191 | 74083 | 74127 | 26 | 26 | 11.56 |
| LCO-G-ArCOIR-B_M0119 | 3333 | 1406 | 9 | 9 | 0 | 0 | 0 |
| LCO-G-ArCOIR-C_M0121 | 235 | 214 | 96525 | 96228 | 37 | 37 | 15.74 |
| LCO-G-ArCOIR-D_M0126 | 424 | 399 | 195 | 198 | 15 | 12 | 2.83 |
| LCO-G-ArCOIR-E_M0129 | 570 | 510 | 129 | 133 | 11 | 11 | 1.93 |
| LCO-G-ArCOIR-F_M0130 | 23721 | 15598 | 214 | 214 | 3 | 3 | 0.01 |
| LCO-G-ArCOIR-G_M0131 | 4727 | 2564 | 288 | 295 | 7 | 7 | 0.15 |
| LCO-G-ArCOIR-H_M0132 | 3491 | 2684 | 236609 | 236766 | 569 | 495 | 14.18 |
| LCO-G-ArCOIR-I_M0134 | 22097 | 13400 | 694 | 686 | 121 | 121 | 0.55 |
| LCO-G-ArCOIR-J_M0136 | 78 | 74 | 433978 | 433085 | 34153 | 26801 | 34360.26 |
| LCO-G-ArCOIR-K_M0143 | 24 | 24 | 834 | 825 | 18 | 18 | 75 |
| LCO-G-ArCOIR-L_M0150 | 14 | 13 | 17676 | 17838 | 4091 | 2567 | 18335.71 |
| LCO-G-ArCOIR-M_M0151 | 1 | 1 | 129 | 126 | 2 | 2 | 200 |
| LCO-G-ArCOIR-N_M0152 | 102 | 93 | 12 | 16 | 0 | 0 | 0 |
| LCO-G-ArCOIR-O_M0156 | 4 | 4 | 6 | 10 | 0 | 0 | 0 |
| LCO-G-ArCOIR-P_M0157 | 91 | 87 | 87 | 98 | 54 | 54 | 59.34 |
| LCO-G-ArCOIR-Q_M0158 | 22 | 21 | 78 | 81 | 10 | 10 | 45.45 |
| LCO-G-ArCOIR-R_M0162 | 26 | 26 | 87 | 87 | 0 | 0 | 0 |
| LCO-G-ArCOIR-S_M163C | 0 | 0 | 18 | 17 | 0 | 0 | NA |
| LCO-G-ArCOIR-T_M0164 | 174869 | 172929 | 23 | 33 | 0 | 0 | 0 |
| LCO-G-ArCOIR-U_M0165 | 47 | 46 | 59 | 65 | 3 | 3 | 6.38 |
| LCO-G-ArCOIR-V_M0166 | 75018 | 74266 | 1369 | 1438 | 387 | 362 | 0.48 |
| LCO-G-ArCOIR-W_M0167 | 0 | 0 | 92 | 133 | 6 | 6 | Inf |
| LCO-G-ArCOIR-X_M0181 | 9 | 9 | 5896 | 5887 | 24 | 24 | 266.67 |
| LCO-G-ArCOIR-Y_M0182 | 0 | 0 | 268 | 258 | 18 | 18 | Inf |
| LCO-G-ArCOIR-Z_M0184 | 97498 | 96915 | 92189 | 92372 | 690 | 684 | 0.7 |
| LCO-H-ArCOIR-A_M0188 | 268 | 235 | 51385 | 51479 | 705 | 687 | 256.34 |
| LCO-H-ArCOIR-B_M0189 | 221 | 159 | 275849 | 276061 | 2060 | 1636 | 740.27 |
| LCO-H-ArCOIR-C_M0190 | 264 | 246 | 19 | 17 | 13 | 13 | 4.92 |
| LCO-H-ArCOIR-D_M0195 | 365 | 347 | 38 | 39 | 3 | 3 | 0.82 |
| LCO-H-ArCOIR-E_M0196 | 243081 | 237334 | 3 | 3 | 0 | 0 | 0 |
| LCO-H-ArCOIR-F_M0201 | 1031 | 777 | 8 | 6 | 0 | 0 | 0 |
| LCO-H-ArCOIR-G_M0202 | 534466 | 435783 | 4 | 4 | 0 | 0 | 0 |
| LCO-H-ArCOIR-H_M0203 | 1004 | 923 | 24 | 32 | 22 | 22 | 2.19 |
| LCO-H-ArCOIR-I_M0208 | 21633 | 18042 | 33 | 34 | 0 | 0 | 0 |
| LCO-H-ArCOIR-J_M0210 | 159 | 148 | 84 | 88 | 4 | 4 | 2.52 |
| LCO-H-ArCOIR-K_M0211 | 22 | 21 | 1 | 4 | 0 | 0 | 0 |
| LCO-H-ArCOIR-L_M0212 | 23 | 21 | 22 | 26 | 0 | 0 | 0 |
| LCO-H-ArCOIR-M_M0219 | 0 | 0 | 8 | 8 | 0 | 0 | NA |
| LCO-H-ArCOIR-N_M0227 | 132 | 105 | 32 | 29 | 0 | 0 | 0 |
| LCO-H-ArCOIR-O_M0228 | 103 | 89 | 13 | 11 | 2 | 2 | 1.94 |
| LCO-H-ArCOIR-P_M0229 | 121 | 117 | 13 | 14 | 0 | 0 | 0 |
| LCO-H-ArCOIR-Q_M0197 | 30 | 25 | 334836 | 334517 | 2916 | 2746 | 9153.33 |
| LCO-H-ArCOIR-R_SphA7 | 49 | 46 | 823 | 846 | 68 | 68 | 138.78 |
| LCO-I-ArCOIR-A_Kik01 | 105 | 90 | 909 | 958 | 18 | 18 | 17.14 |
| LCO-I-ArCOIR-B_Kik02 | 5744 | 1589 | 475525 | 476954 | 588 | 580 | 10.1 |
| LCO-I-ArCOIR-C_Kik03 | 190 | 163 | 545853 | 543435 | 963 | 915 | 481.58 |
| LCO-I-ArCOIR-D_Kik04 | 6188 | 5941 | 312000 | 312604 | 14530 | 7296 | 117.91 |
| LCO-I-ArCOIR-E_Kik05 | 398 | 353 | 634261 | 634738 | 29074 | 23327 | 5861.06 |
| LCO-I-ArCOIR-F_Kik06 | 107751 | 92906 | 397533 | 398442 | 1052 | 978 | 0.91 |
| LCO-I-ArCOIR-G_Kik07 | 57672 | 51835 | 142 | 176 | 48 | 48 | 0.08 |
| LCO-I-ArCOIR-H_Kik08 | 279956 | 277345 | 520751 | 518938 | 1132 | 1113 | 0.4 |
| LCO-I-ArCOIR-I_TAG | 26 | 21 | 104 | 132 | 0 | 0 | 0 |
| LCO-I-ArCOIR-J_TAG | 61 | 54 | 703 | 716 | 4 | 4 | 6.56 |
| LCO-I-ArCOIR-K_TAG | 9 | 8 | 18 | 18 | 0 | 0 | 0 |
| LCO-I-ArCOIR-L_TAG | 13 | 12 | 238321 | 238300 | 62709 | 43760 | 336615.38 |
| LCO-I-ArCOIR-M_TAG | 5 | 5 | 28845 | 28713 | 1145 | 905 | 18100 |
| LCO-I-ArCOIR-N_TAG | 54 | 37 | 591 | 624 | 4 | 4 | 7.41 |
| LCO-I-ArCOIR-O_TAG | 47 | 46 | 64 | 81 | 0 | 0 | 0 |
| LCO-I-ArCOIR-P_TAG | 110 | 105 | 207424 | 207667 | 2220 | 2214 | 2012.73 |
| LCO-I-ArCOIR-Q_TAG | 9 | 9 | 5 | 5 | 0 | 0 | 0 |
| LCO-I-ArCOIR-R_TAG | 34 | 34 | 8727 | 8717 | 234 | 231 | 679.41 |
| LCO-I-ArCOIR-S_TAG | 0 | 0 | 175188 | 174830 | 1743 | 1742 | Inf |
| LCO-I-ArCOIR-T_TAG | 13 | 12 | 147 | 148 | 79 | 75 | 576.92 |
| LCO-I-ArCOIR-U_TAG | 46 | 44 | 3 | 3 | 0 | 0 | 0 |
| LCO-I-ArCOIR-V_TAG | 20 | 18 | 310663 | 310811 | 4046 | 3972 | 19860 |
| LCO-I-ArCOIR-W_TAG | 0 | 0 | 422 | 423 | 156 | 156 | Inf |
| LCO-I-ArCOIR-X_TAG | 0 | 0 | 203 | 199 | 0 | 0 | NA |
| LCO-I-ArCOIR-Y_TAG | 0 | 0 | 290 | 294 | 0 | 0 | NA |
| LCO-I-ArCOIR-Z_TAG | 23 | 21 | 193485 | 193919 | 26 | 26 | 113.04 |

**Table S3**: Information on every sampling site. It shows where the sample was collected, the coordinates (lat and long), the results of Vsearch (Perc, Order, Family, Genus, GenBank), the ASV and OTU found in the sample, the number of reads (count), and drivers (elevation, temperature_max, temperature_min, actual evapotransiration (AET) and canopy cover (cover_code)). For canopy cover (cover_code): 3 is Forest, 12 is Grassland, 13 is Other non-forest formation, 21 is Agricultural mosaic, 33: River, lake and ocean and 68: Other natural non-vegetated area

## Notes

### Competing Interest Statement

The authors have declared no competing interest.

